# SMARTIE: A Machine-Learning approach for investigating RBP-RNA interactions identified by Editing

**DOI:** 10.64898/2026.05.18.726004

**Authors:** Omkar Koppaka, Utham Kumar, Gaurav Ahuja, Rishikesh Yadav, Baskar Bakthavachalu

## Abstract

RNA-binding proteins (RBPs) play important roles in gene regulation. RNA editing-based approaches, such as TRIBE and STAMP, have gained wider use for identifying RNA targets of RBPs. These methods offer advantages over crosslinking-based approaches in terms of experimental simplicity and *in vivo* applicability. However, data analysis methods for these approaches remain underdeveloped, limiting sensitivity, and unbiased target prioritization. To address these limitations, we introduce SMARTIE (Systematic Machine-learning Approach for RBP Targets Identified by Editing), a machine-learning-based framework. SMARTIE robustly identifies and ranks RBP target RNAs from editing data by integrating statistical tests with replicate-aware and confidence-weighted features. Reanalysis of multiple published TRIBE datasets demonstrates the effectiveness of SMARTIE. It recovers targets of RBPs like Ataxin-2, TDP-43, Hrp48, Thor, GPATCH8, dFMRP and NonA. Notably, a model trained on TRIBE data generalizes to STAMP datasets, suggesting that SMARTIE learns universal signatures of editing-based RBP targeting there by enabling more accurate inference for RBP-RNA interactions.

**Graphical Abstract:** 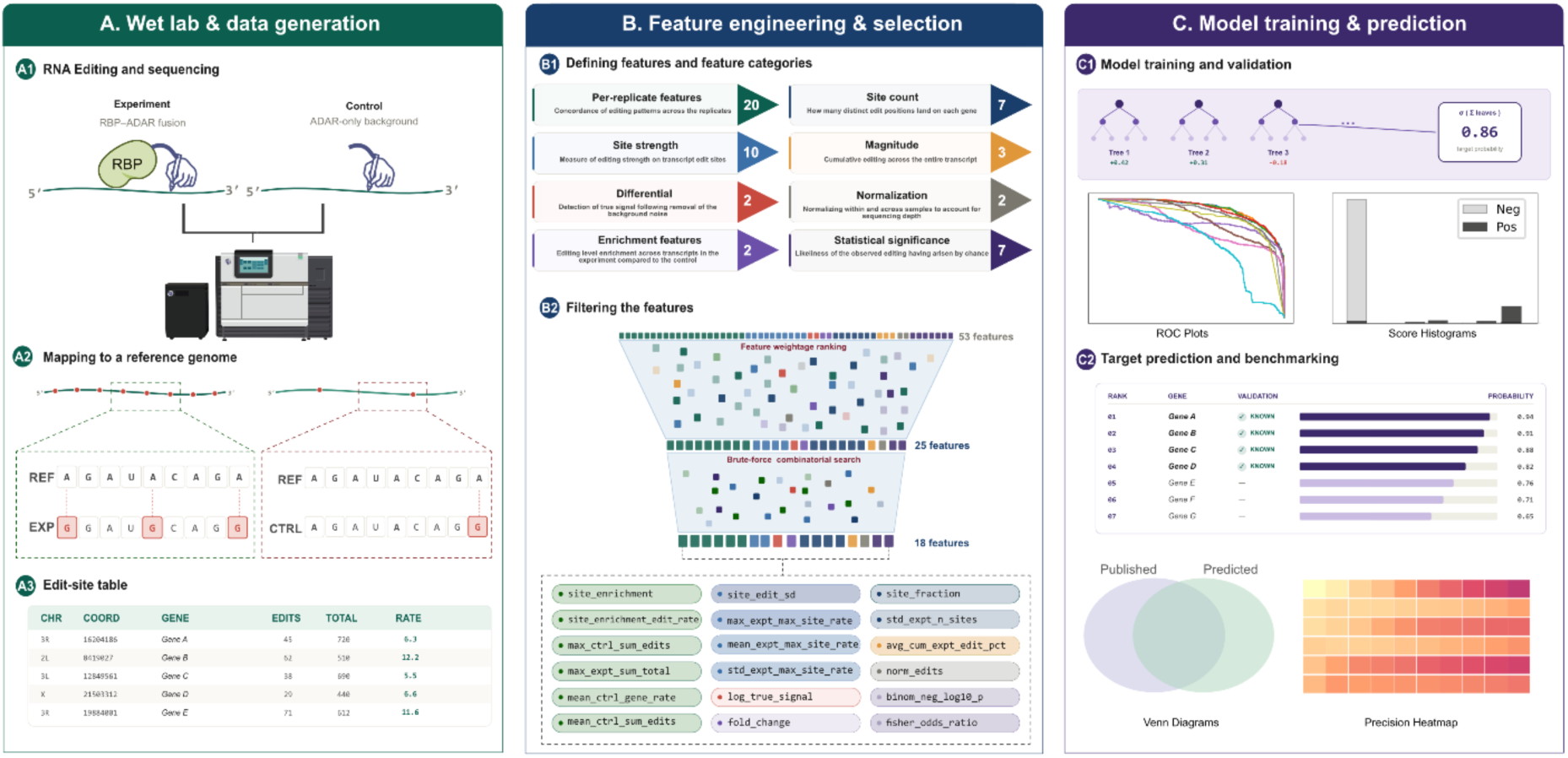

## Introduction

Identifying RNA targets of RBPs (RNA binding proteins) is critical for understanding gene regulation. Techniques for large-scale identification of RNA targets of RBPs are generally categorized as immunoprecipitation-based or RNA editing-based methods. Immunoprecipitation-based methods, such as Crosslinking Immunoprecipitation (CLIP-seq) and its variants, involve pulldown of RNA-RBP complexes followed by RNA purification and identification by sequencing. These methods are widely used for target identification, offering high-resolution mapping of RNA binding sites (Licatalosi et al., 2008; Tenenbaum et al., 2000; Ule et al., 2003; Van Nostrand et al., 2016). However, CLIP-seq and its variants are snapshot techniques that demand large starting material, rely on the availability of specific antibodies, and are influenced by the strength of RBP-RNA interactions. Moreover, with the growing interest in cell-type-specific gene regulation, single-cell transcriptomics has become crucial, highlighting the significant limitation of CLIP-seq (Hafner et al., 2010; Lee & Ule, 2018; Xiang et al., 2024).

RNA editing-based methods provide a promising alternative to overcome these challenges. One of the earliest approaches, known as Targets of RNA binding proteins Identified By Editing (TRIBE), involves fusing the catalytic domain of Adenosine Deaminase Acting on RNA (ADARcd) to the RNA-binding domain (RBD) of the RBP of interest (McMahon et al., 2016). ADARcd edits Adenosine residues by converting them to Inosine (A-to-I), which is read as Guanosine (G) during sequencing (Bass & Weintraub, 1988; Wagner et al., 1989). Similarly, the APOBEC (Apolipoprotein B mRNA Editing enzyme, Catalytic polypeptide-like) enzyme, which converts Cytosine to Uracil (C-to-U), has been utilized in methods such as DART-seq and STAMP (Surveying Targets by APOBEC-Mediated Profiling) (Brannan et al., 2021; Meyer, 2019; Navaratnam et al., 1993). Studies have also explored the combined use of TRIBE and STAMP (Flamand et al., 2022), as well as comparative analyses of their efficiencies (Abruzzi et al., 2023). In addition, these RNA editing-based approaches have been successfully applied to identify RBP targets in single-cells (Brannan et al., 2021; Leeuwen et al., 2022).

Despite their advantages, these methods are constrained by specific limitations. ADARcd and APOBEC introduce edits stochastically near RBP binding sites rather than at precise genomic coordinates. However, due to these properties of the RNA editing enzymes, TRIBE and STAMP are prone to sequence bias and low editing efficiency (Abruzzi et al., 2023; Xu et al., 2018). Recent experimental developments have addressed these issues. The editing efficiency of TRIBE has been improved using an ADARcd mutant (E488Q) (Xu et al., 2018), while STAMP has been optimized using an APOBEC mutant (H122L, D124N) (Jia et al., 2025). However, while experimental protocols have evolved, data analysis pipelines remain underdeveloped for the comprehensive identification of RNA targets. These pipelines, irrespective of the tool used, typically rely on stringent, fixed editing thresholds (e.g., 5–15%) per site and require exact edit coordinate matches across replicates (Abruzzi et al., 2023; Brannan et al., 2021; Leeuwen et al., 2022; Singh et al., 2021; Xu et al., 2018). This leads to the exclusion of many genuine RNA targets and creates significant challenges for experimental reproducibility.

To address these limitations, we developed a transcript-wide, coordinate-independent approach to identify RNA targets of RBPs using a modified method called Systematic Machine-learning Approach for RBP Targets Identified by Editing (SMARTIE). SMARTIE integrates statistical tests with replicate-aware, confidence-weighted features to robustly identify and rank RBP target RNAs from editing data. By capturing the enrichment, consistency, and magnitude of RNA editing, the tool assigns a probabilistic score to each gene candidate. We validated this approach by reanalyzing Ataxin-2 targets in adult *Drosophila* brains (Singh et al., 2021), identifying targets with high confidence and reproducibility. Benchmarking against seven additional RBPs (Ataxin-2, dFMRP, NonA, GPATCH8, TDP-43, Hrp48, and Thor) further confirmed SMARTIE’s effectiveness (Abruzzi et al., 2023; Benbarche et al., 2024; Jin et al., 2020; McMahon et al., 2016; Singh et al., 2021; Xu et al., 2018). Notably, the model demonstrates strong generalization across datasets, indicating that it can be applied to diverse RBPs by learning universal signatures of editing-based targeting, thereby enabling more accurate inference of RBP-RNA interactions.

## Results

### The limitations of the existing TRIBE analysis method

TRIBE is a relatively efficient technique for RNA target identification, as it relies on *in vivo* RNA editing guided by the RBDs of RBPs. Current analysis relies on two primary criteria: (a) the percentage of edits at specific coordinates (Edit percentages or edit rate) and (b) the reproducibility of these edit coordinates across replicates. Edit percentages provide valuable information regarding the strength of RBP binding and ensure reproducibility.

While the current analysis method successfully removes most of the background noise by using the ADARcd only controls and provides a list of targets, a comparison of replicates consistently shows moderate intersection. We used TRIBE data for Ataxin-2 in *Drosophila* brain (Singh et al., 2021) to demonstrate the limitations of TRIBE analysis (Figure. 1a). We propose that the issue arises since existing analysis pipeline does not account for limited stochasticity of ADARcd editing across replicates. The randomness of editing around the RBP binding neighbourhood by ADARcd introduces variability in editing coordinates and alters the edit percentage thresholds of individual sites. Consequently, this leads to poor replicate intersection and the exclusion of some targets as false negatives. Importantly, this limitation cannot be resolved by simple parameter adjustments, such as altering the edit threshold, as this issue would persist across all thresholds.

**Figure. 1:**
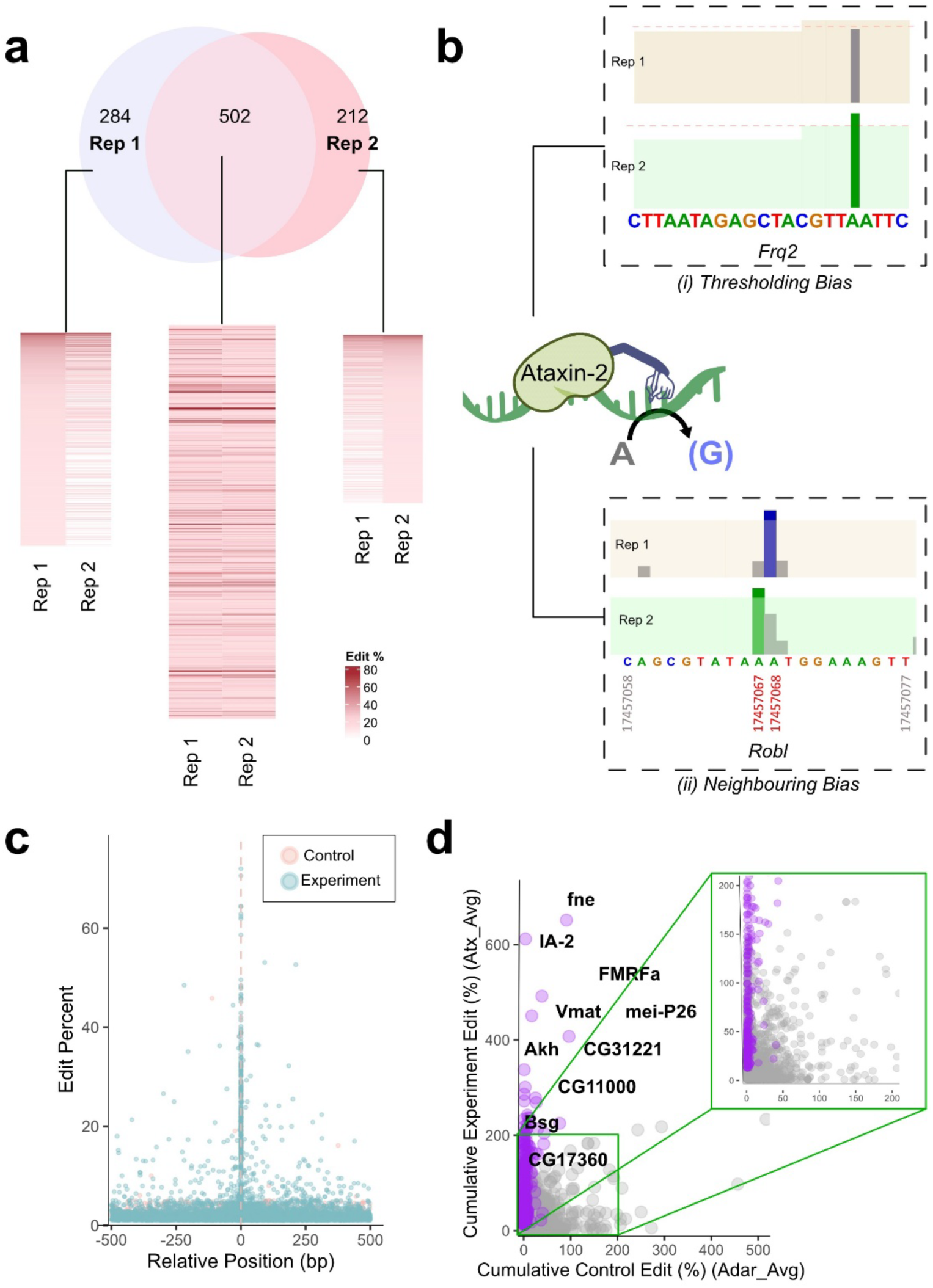
Limitations of existing TRIBE analysis method. (a) Intersection of edits identified in Ataxin-2 TRIBE experimental replicates. The heatmap illustrates differences in edit percentages between replicates. (bi) Edits with respect to threshold for Frq2 gene. (Threshold is represented by red dashed line) (bii) Clustered edits identified in Robl gene near RBP binding sites. (c) A global edit distribution plot for Ataxin-2 targets around the primary edit coordinates. (d) Scatter plot comparing cumulative editing percentages for experimental (Y-axis) and control (X-axis) samples. The TRIBE targets identified using existing analysis method are highlighted in purple while the zoomed section shows presence of multiple genes having similar parameters as the target genes.

Manual evaluation of specific genes confirmed this threshold bias, with some genes narrowly missing the cutoff in one replicate and being excluded from the final target list (Figure. 1b-i). In other cases, where adenosines are clustered near the RBP binding sites, ADARcd edited adjacent adenosines across replicates, introducing coordinate variability (Figure. 1b-ii). When a global edit distribution plot was generated by plotting the edit percentages around the primary edit, a distinct edit distribution is observed around the primary edit peak, consistent with a localized ADARcd editing footprint. (Figure. 1c).

As the editing near the RBP binding site is stochastic, the RBP-ADARcd can introduce edits within an ∼500 bp window (Xu et al., 2018). However, the total edits on a target mRNA are anticipated to be more noticeable in the experiment compared to the control. Consequently, calculating the cumulative edit percentage for each transcript and comparing the results across experimental and control conditions may offer a more reliable indicator of target identification. To test this hypothesis, all the edits above 5% on an RNA were added to calculate a cumulative edit score for each gene. This process was repeated for both experimental and control samples. Following this, cumulative edit percentages were averaged across replicates for every gene (Figure. S1a). A scatter plot comparing average experimental cumulative edit percentage on Y-axis against average control cumulative edit percentage on X-axis showed that potential targets cluster near the Y-axis above the intersection (Figure. 1d). Genes in the bottom-left quadrant, although following the same trend as the bonafide targets (shown with purple dots), were not identified by the conventional TRIBE analysis (Figure. 1d, zoomed). Overall, these findings highlight the critical limitations of coordinate-only approach in the current TRIBE analysis pipeline and support the need for further refinement.

### Transcript-level editing signatures improves RNA target identification

To improve the efficiency of target identification, we calculated a fold change value by dividing the experimental average cumulative edit percent with the corresponding value in the control. The genes with cumulative edits below 15% in the experimental dataset were excluded to improve specificity. Additionally, genes with zero cumulative editing in control samples were assigned a pseudocount of 1 to enable fold-change calculations and avoid division by zero. Fold changes were then calculated as the ratio of cumulative edit percentages in experimental versus control samples (Figure. S1a).

Using this approach, we identified 1,343 genes that had FC≥2 as Ataxin-2 targets in the *Drosophila* brain. Approximately 88% of targets originally reported overlapped with the targets identified using the cumulative edit fold change calculations (Singh et al., 2021) (Figure. S1b). More importantly, the additional targets identified using transcript-level editing signatures had the same characteristics as Ataxin-2 targets, such as their 3’UTR specificity (Figure. S1c). These findings demonstrate that combining transcript-level aggregation overcomes key limitations of coordinate-dependent analyses, providing a reliable framework for RNA target identification.

However, the use of transcript-level editing fold-change alone has its limitations. Like the existing coordinate-based approach, it relies on arbitrary thresholds, making it inconsistent across laboratories. Furthermore, a fixed fold-change threshold of ≥2, risks excluding genuine targets of weak RBPs with a limited binding sites. To address these issues, we developed a machine learning model that incorporates transcript-level cumulative edit percentages and other gene level parameters of TRIBE, to train a generalized model capable of accurately predicting RNA targets from any TRIBE experiments.

### Reanalysis of published TRIBE datasets with a standardized preprocessing for machine-learning model development

TRIBE data for RBPs, including Ataxin-2, TDP-43, Hrp48, Thor, GPATCH8, dFMRP, and NonA was used for training and testing the machine-learning model in addition to STAMP data for TDP-43 (Abruzzi et al., 2023; Benbarche et al., 2024; Jin et al., 2020; McMahon et al., 2016; Singh et al., 2021; Xu et al., 2018). These samples were selected because the experiments were conducted in different tissues or model organisms to develop a generalizable model that is not biased by sample characteristics. To ensure consistency, we reanalyzed all the TRIBE and STAMP data with a standardized preprocessing protocol.

In summary, the RNA edits coordinates were identified using HyperTRIBE software (Table. S1). Appropriate changes were made in the scripts to account for the C-to-U editing in the STAMP data. Corresponding genomic DNA was used to compare RNA sequences and identify edits. For conventional TRIBE and STAMP analysis, the coordinates with at least 20 sequencing reads and 15% editing were used for further downstream analysis (Figure. S2a). To verify pipeline reproducibility, the reanalyzed target lists were compared against the originally published targets (Figure. S2b, Table. S2). While we find a significant number of overlapping targets to the published list, we also observed some differences, which could be attributed to differences in the mapping tools and genome versions used. For instance, the original analysis for GPATCH8 was done using hg19 as the reference genome while we used hg38 which is the latest available reference genome for human samples (Benbarche et al., 2024).

### Building a machine learning model for TRIBE

TRIBE data, being tabular and feature-rich, is well suited to machine learning algorithms capable of classifying and ranking RNA targets. For this purpose, XGBoost (eXtreme Gradient Boosting) (Chen & Guestrin, 2016) and Random Forest (Ho, 1995) were selected as candidate algorithms, given their demonstrated success in prior studies on structured biological and omics data with comparable characteristics (Abbas et al., 2023; Kim et al., 2016; Nguyen et al., 2022; St Laurent et al., 2013). The core analytical pipeline consisted of three stages: extraction of raw edit files from TRIBE experiments, construction of informative features from these files, and training the model to generate ranked predictions with associated probabilities. Predicted targets were subsequently validated by comparison against the standardized target lists (Figure. 2a). Nine core features were constructed based on biological reasoning and their demonstrated ability to discriminate experimental from control editing patterns (Figure. S3).

**Figure. 2:**
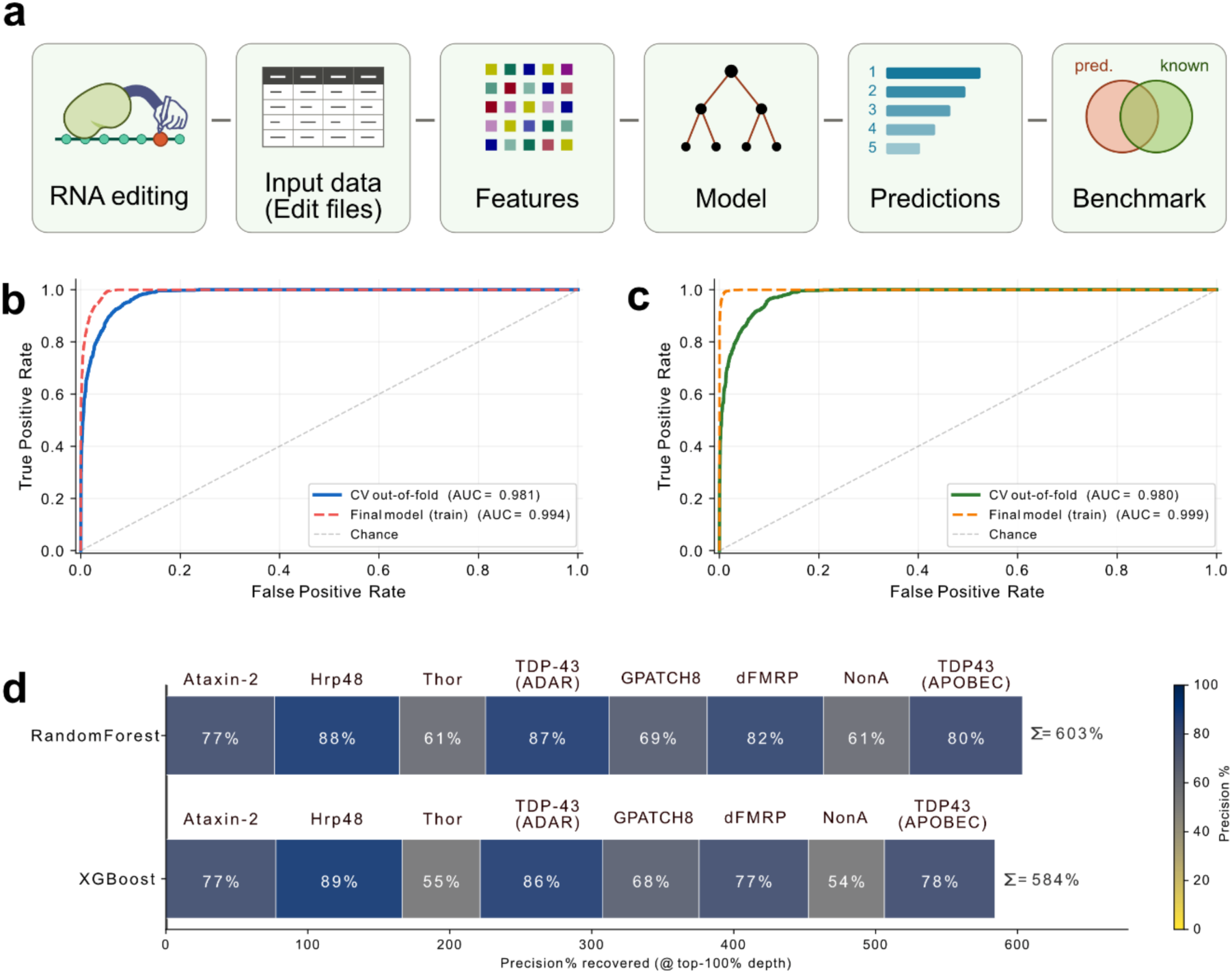
Constructing a machine-learning model for TRIBE. (a) Illustration of workflow for constructing a machine-learning model using TRIBE data. (b) ROC plots for evaluation of Random Forest and (c) XGBoost efficacy. (d) Precision heatmap for Random Forest and XGBoost in identifying the RBP targets.

NonA TRIBE data from *Drosophila* brain (GSE116400) was used as the training dataset and supplied independently to both Random Forest and XGBoost. Both models showed excellent discriminative ability, achieving AUC values of 0.981 and 0.980 for Random Forest (Figure. 2b) and XGBoost (Figure. 2c) respectively. To evaluate how well each model recovers known RBP targets on independent held-out data, performance was quantified using a ranked overlap analysis. Seven different RBPs from 8 datasets were chosen for this purpose (Table.1). For each RBP, the reanalysed target list was compared with an equivalent number of top-ranked SMARTIE predictions, and the percentage overlap was computed. For example, in the analysis of Hrp48 HyperTRIBE data, the target set predicted with existing method comprised 2,842 genes; to evaluate performance, the top 2,842 SMARTIE-ranked genes were intersected with this full list and the resulting overlap fraction calculated.

**Table. 1:**
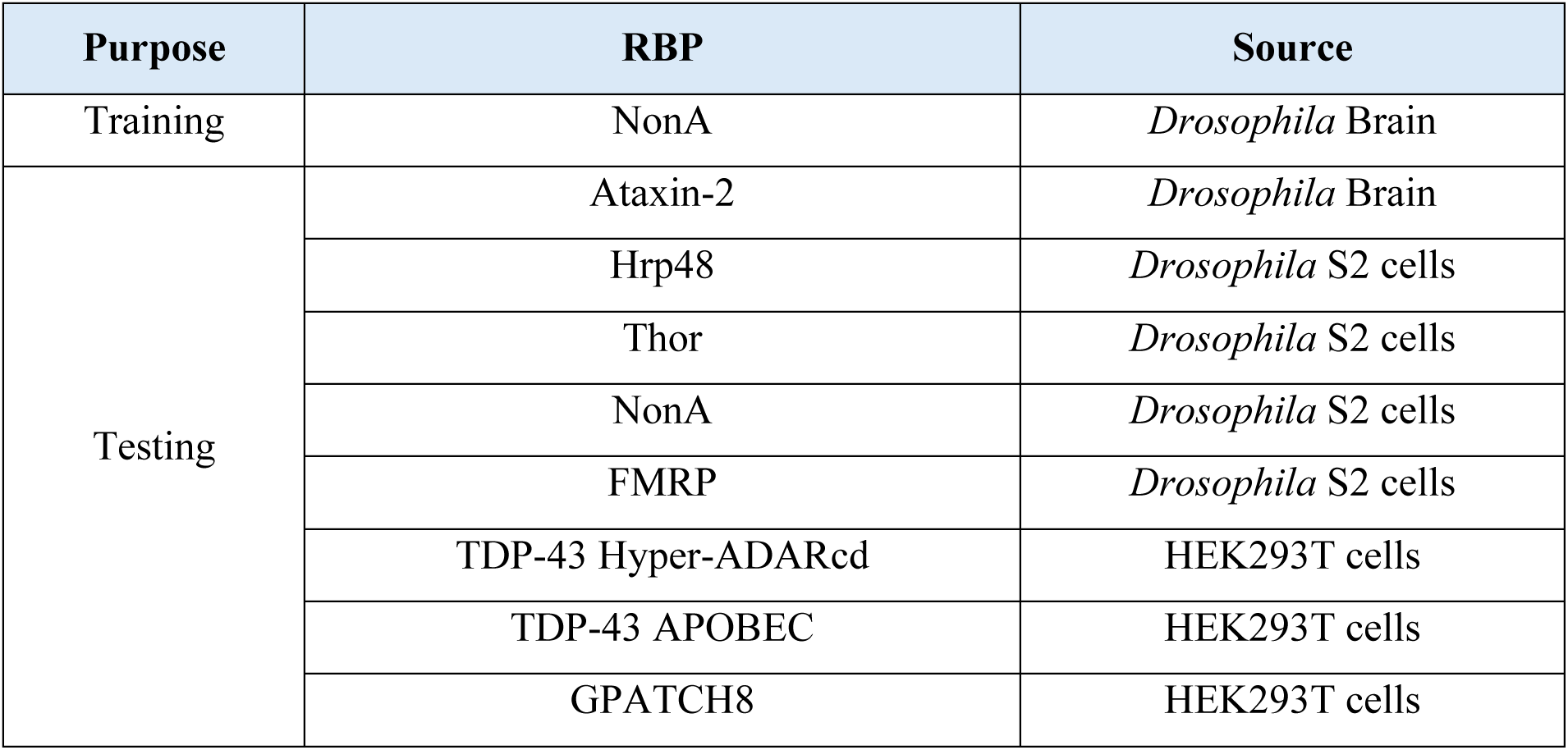
List of Training and Testing TRIBE Datasets.

Both Random Forest and XGBoost performed strongly for RBPs with large target repertoires, such as Hrp48, TDP-43, and Ataxin-2. However, both models showed reduced performance for RBPs with comparatively fewer targets, including NonA, Thor, and GPATCH8 (Figure. 2d). While the core feature set provides a solid foundation for target prediction, these results indicated that further feature optimization would be necessary to improve generalizability across RBPs with diverse binding profiles.

### Feature selection and optimization for machine-learning analysis of TRIBE data

The optimal performance of a machine-learning model hinges on the judicious selection of features. The original nine core features, which were designed to encapsulate the statistical properties of TRIBE editing data and span eight broad categories (Figure. 3a). In addition, a set of 44 additional features (Table. S3) were constructed to capture the key patterns in TRIBE editing data. The optimization was performed using TRIBE data for Ataxin-2, TDP-43, Hrp48, Thor, GPATCH8, dFMRP, and NonA, to find a set of generalized features.

**Figure. 3:**
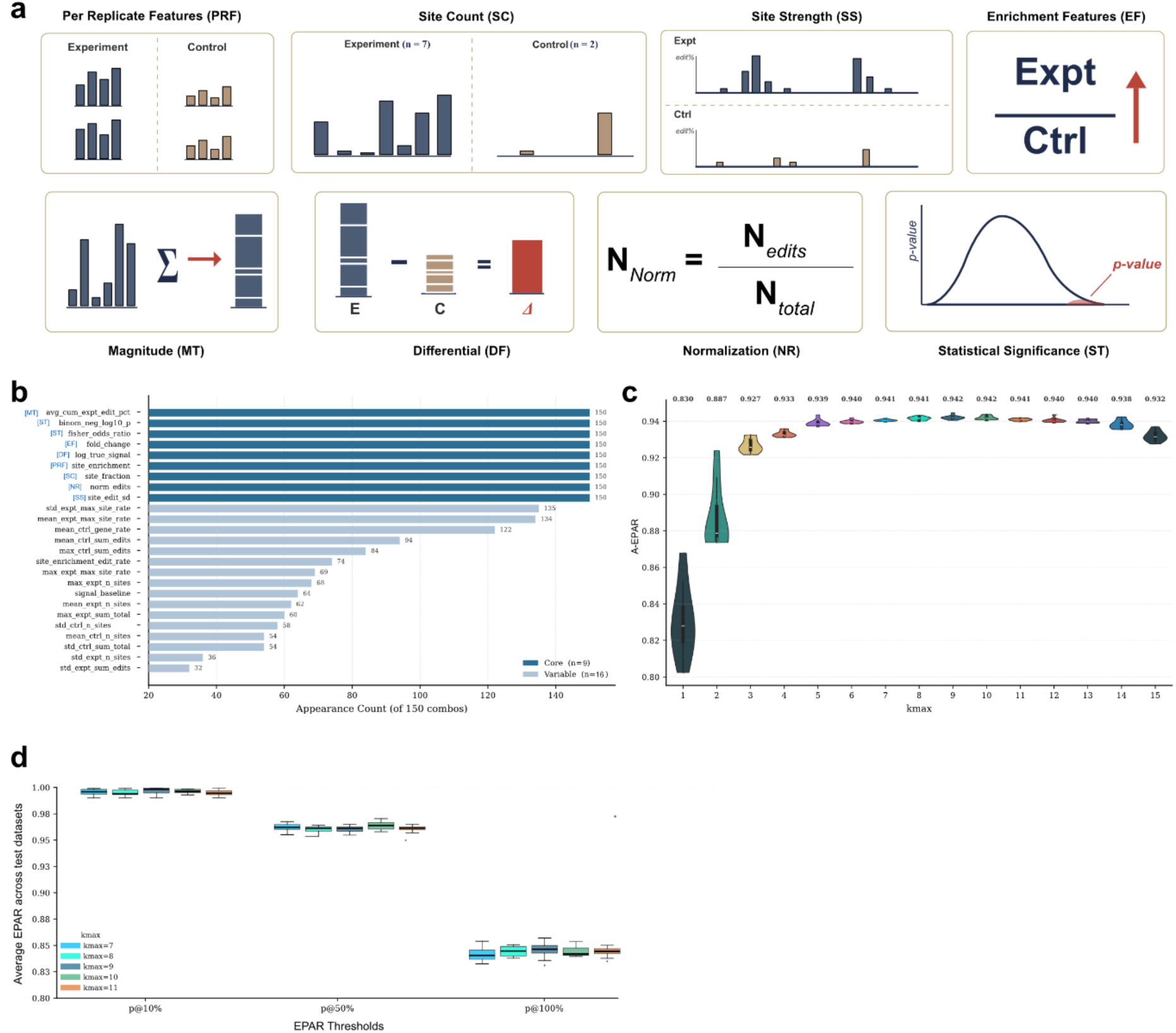
Feature selection and optimization using brute-force method. (a) Schematic overview of the eight feature categories used to capture TRIBE editing patterns. (b) Bar plot depicting the frequency of appearance of each feature across the top 10 combinations at each k-max value from 1 to 15, highlighting the most consistently informative features. (c) Violin plots show the distribution of A-EPAR scores across the top 10 feature combinations at each k-max from 1 to 15, illustrating how overall model performance evolves with increasing feature set size. (d)Box plots comparing combination-level EPAR across classification thresholds of 10%, 50%, and 100% for k-max values of 7 to 11, demonstrating performance stability around the optimal feature set size.

The most informative additional features were selected using a brute-force combinatorial feature selection strategy using SMARTIE. Exhaustive enumeration across all 53 constructed features (9 core + 44 additional features) was computationally prohibitive. For instance, at k-max = 5, the search space exceeds one million combinations (Table. 2). So, we first conducted a discovery run at k-max = 4 (∼100,000 combinations), then ranked features by their frequency of occurrence in high-performing subsets. The top 25 most recurrent features, encompassing the nine core features, were carried forward for evaluation on an XGBoost and Random forest models.

**Table. 2:** Number of feature combinations evaluated perk-max for the two-stage brute force selection strategy.

| **k-max** | **44 variable features** | **16 variable features** |
| --- | --- | --- |
| 1 | 44 | 16 |
| 2 | 946 | 120 |
| 3 | 13,244 | 560 |
| 4 | 135,751 | 1,820 |
| 5 | 1,086,008 | 4,368 |
| 6 | 7,059,052 | 8,008 |
| 7 | 38,320,568 | 11,440 |
| 8 | 177,232,627 | 12,870 |
| 9 | 708,930,508 | 11,440 |
| 10 | 2,481,256,778 | 8,008 |
| >10 | 845,671,815,201 | 6,885 |

With the reduced 25-feature set, the brute-force selection was repeated (Figure. 3b). The substantially smaller search space (∼65,000 total combinations) permitted more thorough exploration (Table. 2). To evaluate model performance during brute-force feature selection, we defined a metric termed the Equivalent Precision Against Reference (EPAR). For each RBP, the reanalyzed target list was partitioned into deciles of 10% ranked bins. For each decile, an equivalent number of top-ranked SMARTIE predictions was extracted and intersected with the complete reanalyzed target list, and the percentage overlap was computed. For example, in the analysis of Hrp48 HyperTRIBE data, the reanalyzed target set comprised 2,842 genes; to evaluate performance at the 10% threshold, the top 284 SMARTIE-ranked genes were intersected with this full list and the resulting overlap fraction calculated. This procedure was repeated across all deciles and RBPs, and decile-level overlaps were averaged across datasets to generate a single summary EPAR value per feature combination, which served as the primary metric for evaluating model performance throughout the brute-force selection process. To assess performance as a function of kmax, EPAR values were averaged across all datasets and all decile thresholds.

Average EPAR (A-EPAR), which averages EPAR across all datasets and thresholds, rose from 0.83 at kmax = 1 to a peak of 0.942 at kmax = 9–10, followed by a negligible decline to 0.932 at kmax = 15 (Figure. 3c), indicating that performance plateaus beyond kmax = 9 with no meaningful gain from adding further variable features. To confirm that this result was not driven by any particular classification threshold, performance assessment was carried out across thresholds 10%. 50% and 100%. For k-max values of 7 to 11; EPAR remained consistently high across this range (0.941–0.942), and the advantage of the optimal feature set was maintained at every threshold examined (Figure. 3d). Given that k-max = 9 yielded both the highest overall precision and the most stable per-threshold performance, an optimized SMARTIE model was adopted, incorporating 18 optimized features (9 core + 9 variable).

### XGBoost most efficiently prioritizes features for SMARTIE

Next, we compared XGBoost (Chen & Guestrin, 2016) with several other classification algorithms (Gradient Boosting (Friedman, 2001), Naïve Bayes (Watson, 2001), SVM (Vapnik, 1997), Logistic Regression (Cramer, 2002), LightGBM (Ke et al., 2017), Random Forest (Ho, 1995), Linear Regression (Kuchibhotla et al., 2019), K-Nearest Neighbor (Seidl, 2009) and Decision trees (Rokach & Maimon, 2005) against the 18 finalized features to identify the most suitable algorithm for TRIBE (Figure. 4a). The models were trained on the NonA brain dataset and evaluated using precision-recall metrics. Gradient Boosting achieved the highest average precision (0.93), followed closely by XGBoost and LightGBM (both 0.92), while Naïve Bayes performed poorly (0.689) (Figure. 4b). Train-versus-cross-validation ROC curves confirmed that all models fit the training data well, and most, with the exception of Naïve Bayes, SVM, and Logistic Regression, were able to discriminate targets from non-targets effectively (Figure. 4c).

**Figure. 4:**
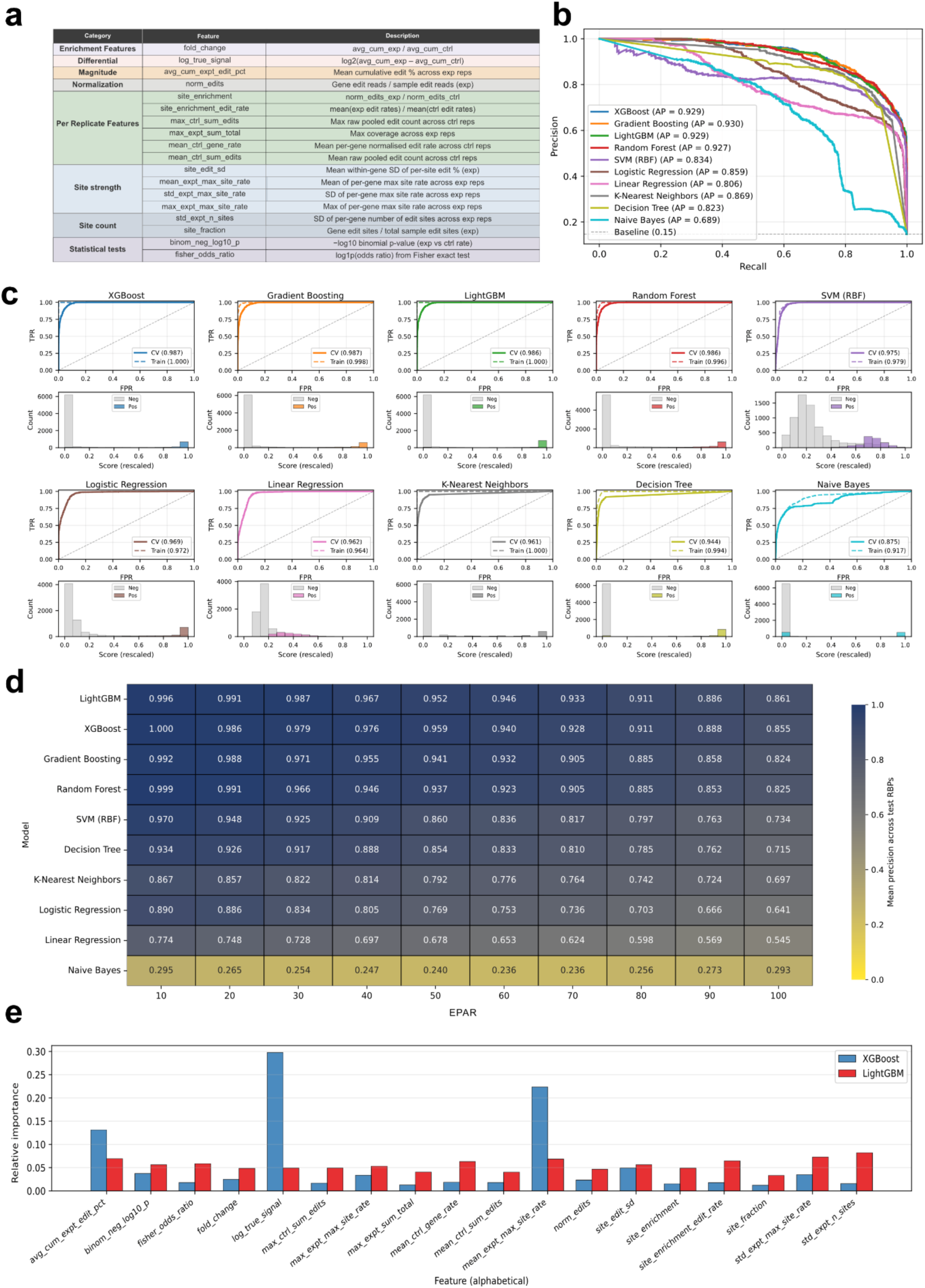
Comparative evaluation of classification models identifies XGBoost as the optimal algorithm for SMARTIE. (a) Description for each of the 18 features selected for training SMARTIE. (b) ROC curves for all ten candidate models evaluated on the training dataset, illustrating their ability to discriminate RBP targets from non-targets. (c) Paired comparison of training and cross-validation ROC curves (top) alongside the distribution of predicted scores for targets and non-targets (bottom), used jointly to assess model overfitting and discriminative capacity. (d) Heatmap depicting averaged EPAR value for all the test datasets at each decile threshold across all held out testing datasets, enabling quantitative comparison of target recovery performance across models. (e) Feature importance plots for XGBoost and LightGBM, illustrating the differential feature prioritization strategies employed by each model.

High precision-recall and ROC values, however, raise the concern of overfitting, and performance on training data alone cannot be taken as evidence of real-world generalizability. So, the models were also evaluated on other RBPs like Ataxin-2, TDP-43, Hrp48, Thor, GPATCH8, dFMRP. Model performance was quantified using a ranked overlap analysis and visualized as a heatmap.

LightGBM and XGBoost emerged as top-performing models, with marginal differences followed by Gradient Boosting and Random Forest based on the EPAR values (Figure. 4d). To distinguish between them, we examined how each model allocated feature importance. Notably, XGBoost utilized variable weights for features, LightGBM distributed importance more diffusely, with no dominant choices (Figure. 4e). Considering that true TRIBE targets are likely to comprise a limited subset of genes and to carry concentrated signal, we reasoned that a model capable of focusing on the most informative features would be better suitable for this task. Therefore, XGBoost was chosen as the final algorithm for SMARTIE.

### SMARTIE accurately identifies TRIBE and STAMP targets

SMARTIE predictions for all RBPs were independently calculated and compared with the targets from existing analysis that had been partitioned into deciles. For each decile, an equal number of top-ranked SMARTIE predictions were extracted, intersected, and the percentage overlap was calculated and presented as a heat map (Figure. 5a and Table. S4). SMARTIE achieved a near perfect EPAR at the fifth decile of its ranked predictions across all the 8 test datasets, 100% for Hrp48, TDP-43, and dFMRP; 99% for Ataxin-2 and 97% for Thor, and 87% for S2 cell NonA, with a minimum EPAR of 83% for GPATCH8. Even at the tenth decile, SMARTIE predictions overlapped with an average of 85.5% of the complete reanalyzed target lists across RBPs, reaching a maximum EPAR of 94% for the Hrp48 datasets.

**Figure 5:**
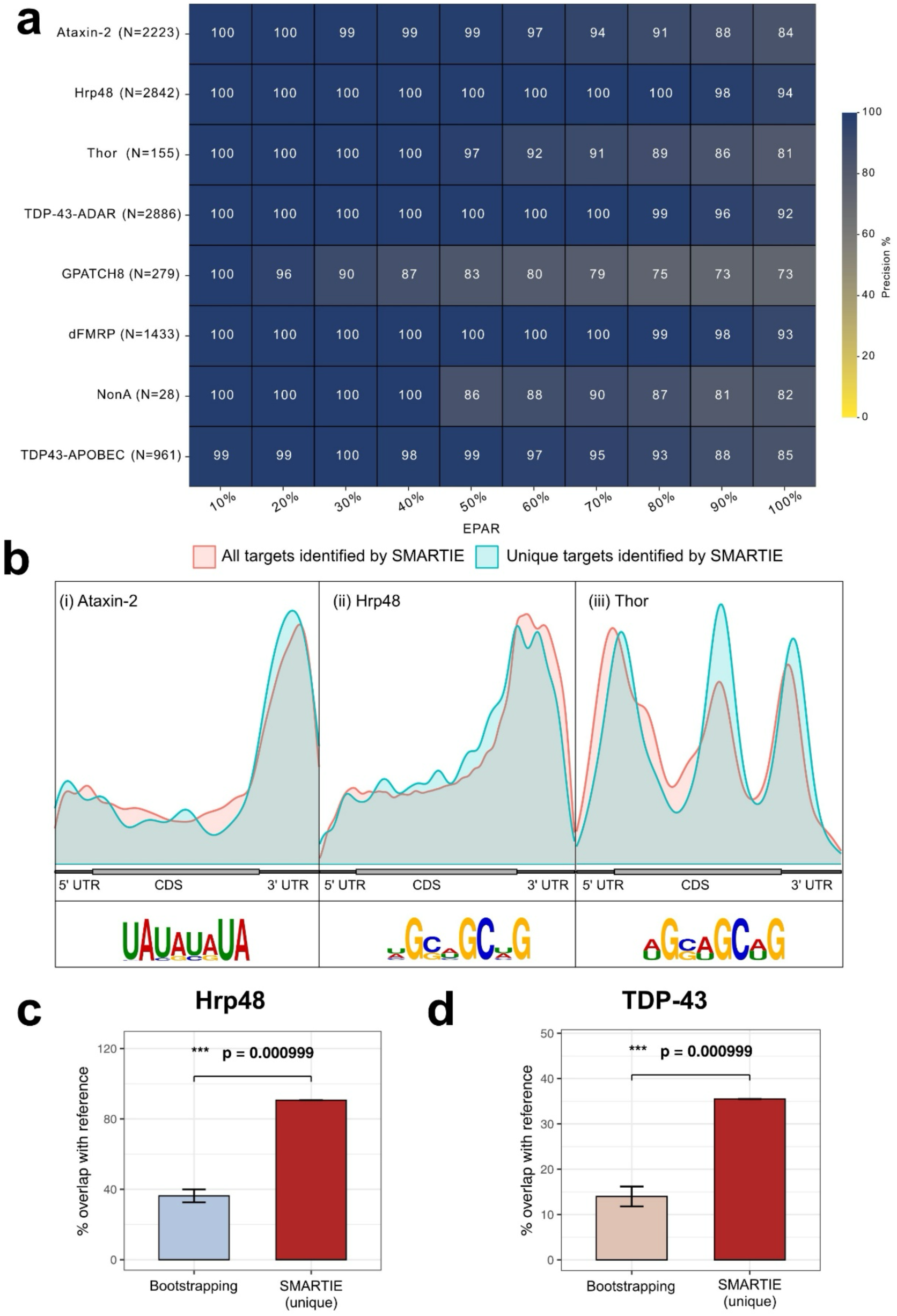
Benchmarking SMARTIE using TRIBE and STAMP datasets. (a) A heatmap for decile-based analysis of SMARTIE-predicted target intersected with the reanalysed list of targets. (b) Metaplot and motif for (i) Ataxin-2, (ii) Hrp48, and (iii) Thor. Overlap of SMARTIE-unique targets for Hrp48 (c) and TDP-43 (d) with CLIP datasets (red) versus a bootstrapped random-sampling baseline (1,000 iterations; blue/pink). ***p = 0.000999.

To study the generalizability of SMARTIE, it was tested on TDP43 STAMP which employs APOBEC enzyme mediated converts cytosine to uracil (C-to-U) editing. While this editing signature differs from TRIBE A-to-I editing, SMARTIE successfully identified TDP43 STAMP targets that were comparable to the TRIBE predictions (Figure. 5a).

Meta-analysis, including motif extraction and RBP binding positional preference mapping was performed for SMARTIE-predicted targets using editing-site coordinates for the target genes. RBP-specific binding characteristics were observed. Ataxin-2 targets showed strong enrichment in 3′ UTRs and were associated with AU-rich motifs (Figure. 5b-i), aligning with its known binding preferences (Singh et al., 2021; Yokoshi et al., 2014). Hrp48 targets were enriched in 3′ UTRs with CG-rich sequence (Figure. 5b-ii), while Thor targets showed preferential binding in 3′ and 5′ UTRs associated with CG and CU motifs (Figure. 5b-iii). The targets that were uniquely identified by SMARTIE but not by conventional analysis were extracted and subjected to motif analysis and metaplot. The SMARTIE-unique targets accurately matched the predicted characteristics for all three RBPs.

To further validate SMARTIE, we obtained independent CLIP data from the POSTAR3 database for TDP-43 (Zhao et al., 2022) and from McMahon et al., 2016 for Hrp48. SMARTIE-unique targets exhibited a remarkable 91% overlap with the CLIP data for Hrp48 and a 35.5% overlap with TDP-43. Notably, this overlap is approximately 2.5-fold higher than the randomly selected equivalent number of Drosophila genes used for bootstrapping the background in each case (Figure. 5c and d).

Collectively, these results demonstrate that SMARTIE learns transferable statistical signatures of bonafide RBP targets across RNA editing-based experimental platforms. The faithful recovery of these RBP-specific interaction signatures across multiple proteins confirms that SMARTIE predictions are not only statistically robust but also biologically meaningful.

## Discussion

TRIBE and STAMP are emerging techniques with broad applications in RNA target identification. However, their effective use requires analytical methods specifically tailored to the unique biochemical and statistical properties of RNA editing enzymes. Conventional TRIBE analysis identifies editing events by comparing experimental samples to a reference dataset at specific genomic coordinates to detect nucleotide conversions. A fundamental limitation of coordinate-dependent analysis comes from the stochastic editing by ADAR or APOBEC near the RBP binding site that leaves variability in replicate experiments.

Use of target transcript level aggregate editing levels overcomes this key limitation. We therefore recognized the need for an unbiased, coordinate-independent method grounded in a principled statistical framework. This motivated the development of SMARTIE, a post-processing analytical tool designed to enable reproducible and comprehensive RNA target identification from RNA editing data.

Machine learning model performances depend on feature design and algorithm selection. Therefore, a list of 53 features spanning eight broad categories were constructed that collectively capture the key dimensions of TRIBE editing data (Figure. 2a, Table. S3). While ensuring a set of biologically motivated core features, a brute-force combinatorial selection was applied to finalize feature list to evaluate using both Random Forest (Figure. S4) and XGBoost algorithms (Figure. 3) with NonA brain TRIBE as training data. XGBoost was ultimately selected as the algorithm underlying SMARTIE due to its robustness, accuracy and the ability to assign mixed feature weights. Using SMARTIE, we identified targets for Ataxin-2, Hrp48, Thor, TDP-43, GPATCH8, dFMRP and NonA.

Unlike earlier TRIBE and STAMP pipelines that produce fixed binary target lists, SMARTIE generates a continuous ranked output, allowing users to define confidence thresholds appropriate to the biological context and binding properties of the RBP under investigation. Genes ranked highly by SMARTIE, showed strong concordance with the target datasets for all the tested proteins demonstrating robustness across diverse RBPs and experimental systems (Figure. 5). These results consistently demonstrate high predictive accuracy across diverse RBPs and experimental systems, consistent with the earlier observation that high-confidence targets are reliably identified by conventional methods. The marginal reduction in recovery for S2 cell NonA and GPATCH8 likely reflects the inherently limited target repertoire of these RBPs rather than a failure of the model. Notably, SMARTIE performed well even on single-replicate TRIBE datasets, yielding target rankings highly comparable to those from multi-replicate analyses (Figure. S6 and Table S5). This represents a meaningful practical advantage, as it suggests that reliable target identification may be achievable at substantially reduced sequencing cost. Furthermore, SMARTIE’s performance was not constrained by the scale of RBP binding: it accurately recovered targets for highly promiscuous RBPs such as Hrp48 and TDP-43, which each bind thousands of targets, as well as for selective binders such as NonA, which has fewer than 30 known targets in S2 cells. Importantly, SMARTIE-unique targets for Hrp48 and TDP-43 showed significant overlap with independent CLIP data, well above a bootstrapped random baseline, confirming that these are genuine RBP targets (McMahon et al., 2016; Zhao et al., 2022).

Although SMARTIE was trained on TRIBE datasets, we hypothesized that its statistical framework would generalize to STAMP data, given that both methods rely on enzymatic RNA editing and differ primarily in base conversion chemistry (A-to-I versus C-to-U). Consistent with this hypothesis, SMARTIE accurately recapitulated known targets from STAMP datasets, indicating that it captures features of RNA editing that extend beyond the specific editing modality. Importantly, while SMARTIE was trained on a *Drosophila* dataset, it successfully identified targets of TDP-43 and GPATCH8 expressed in human cells using HyperTRIBE. This cross-species generalization suggests that the statistical signatures of RNA editing are broadly conserved. Building on this insight, future efforts will focus on assembling shared control resources to reduce experimental burden, and on developing a web-based interface to make SMARTIE accessible to researchers without specialist bioinformatics expertise.

Several limitations merit acknowledgment. The current model was trained primarily on the NonA brain dataset, and the relatively limited availability of STAMP data precludes comprehensive benchmarking in that context. As with all machine learning approaches, expanding the diversity and size of the training data will further improve performance and generalizability. To facilitate this, SMARTIE is freely available and fully customizable, allowing users to retrain the model or adapt feature sets to new experimental systems. Although SMARTIE was originally developed as a command-line tool, we additionally provide a locally hosted graphical user interface (GUI) that accepts raw experimental and control files. Beyond target prediction, the GUI also supports training and cross-testing of custom models and datasets, allowing users to adapt SMARTIE to new RBPs and experimental systems without programming expertise (Figure. S7). In parallel, we are developing ServerTRIBE, a server-optimized variant of the TRIBE analysis workflow that, together with SMARTIE, will streamline the full analysis pipeline and make TRIBE data analysis accessible to non-bioinformaticians.

In conclusion, SMARTIE provides a robust and flexible analytical framework for RNA target identification using editing-based methodologies. By moving beyond the constraints of coordinate-dependent analyses, it improves sensitivity, reproducibility, and interpretability while substantially reducing false negatives. These advances establish SMARTIE as a broadly applicable platform for RNA-binding protein target discovery across editing-based experimental approaches.

## Methods

### Data retrieval and processing

The NGS data for *Drosophila* and mammalian TRIBE and STAMP experiments were obtained from NCBI GEO database repository and listed in Table. S1. ADARcd-only or APOBEC-only expressing samples were used as “control” and the RBP-ADARcd and RBP-APOBEC fusion were the “experimental” samples. At least two replicates were analysed for each of the conditions.

The FASTQ files were downloaded using the SRA toolkit and the reads were quality tested using FASTQC. Where required, the reads were trimmed using trimmomatic and mapped to appropriate genomes (dm6 for *Drosophila melanogaster* and hg38 for *Homo sapiens*), using STAR. Genomic DNA sequences (gDNA) for human and *Drosophila* were downloaded from PRJNA565658 and GSE37232 respectively. The resulting SAM output files were sorted, de-deduplicated and loaded as a table into a MySQL database. The RNA tables of all samples were compared to its appropriate reference gDNA table to identify edit coordinates as described in McMahon et al., 2016. Fly gDNA was used as reference for Ataxin-2 and NonA fly brain data, S2 cell gDNA was used as reference for Hrp48, NonA-S2, dFMRP & Thor. HEK293T gDNA was used as reference for TDP-43 and GPATCH8. The last step of TRIBE pipeline was modified to detect C-to-U transitions for STAMP analysis.

### TRIBE analysis

Edit percentages were calculated by dividing the total number of A-to-G(I) or C-to-T(U) conversions observed at a specific coordinate by the total number of reads observed at that coordinate. All edits below the 15% threshold were excluded. Additionally, coordinates with sequencing depth less than 20 reads were not considered for analysis. For data from paired-end sequencing, the individual read pairs were processed independently and merged by appending unique edits of Read pair 2 to Read pair 1 before further processing. The control replicates were further merged and subtracted from experimental edits to obtain the final experimental edits. At least two replicates (replicate 1 and replicate 2) were used for each experimental condition, and the final target genes were identified by intersecting the identical genes between the replicates. Further, any edits with an edit frequency ≥95% were excluded from the final list.

### Edit distribution plot

For each coordinate identified using TRIBE, the editing position was designated as the 0 bp reference position. The edit distribution around −50 and +50 bp from the reference position was extracted from the raw edit file ensuring at least 20 sequencing reads. Subsequently, an edit distribution plot/metaplot was generated by plotting the edit coordinate against the edit percentages for targets in experimental and control samples.

### TRIBE target identification using cumulative edits

TRIBE analysis was performed as described earlier (“TRIBE Analysis” section) with the exception of using 5% threshold to generate an edit threshold file. The background edits were subtracted from experimental and control samples. Further, edits with frequency ≥95% were excluded for all samples. The cumulative edit percentage was calculated for each gene by summing the edit percentages of all coordinates for that gene. This created a table with gene names and corresponding cumulative edit percentages. After averaging the cumulative edits for replicates, the genes with cumulative edit percentage of <15% in experimental samples were removed from experiment and control samples. Further, in control samples, rows containing edit percentage zeros were replaced with a value of 1. The average cumulative edits for the experimental condition were divided by the corresponding values for the control condition to calculate the fold-change for each gene. Genes with a fold-change ≥2 were considered as potential targets and the rest were classified as non-targets.

### Machine-Learning for RNA target identification using RNA editing

SMARTIE is a machine learning pipeline implemented in Python and organized into four sequential stages: data loading, feature construction, model training, and prediction. Detailed documentation and all associated scripts are publicly available at https://github.com/toolsmnl/SMARTIE.

#### Data preprocessing

All count columns are converted to numeric values, with non-numeric entries coerced to NaN. Rows with missing values in either count column or the gene name column are subsequently dropped. Edit percentage at each site is then calculated by dividing the edit count by the total count and multiplying by 100. Gene names are standardized by conversion to lowercase with leading and trailing whitespace removed, preventing erroneous duplicate gene entries arising from formatting inconsistencies across datasets. Finally, sites with a total read coverage below 20 are excluded prior to model training and testing using the –min-reads flag.

#### Feature construction and selection

An initial pool of 53 candidate features was constructed based on biological reasoning, of which nine were designated as core features on the basis of their demonstrated discriminative ability (Figure. S3). The remaining 44 features were treated as variable features subject to selection. To identify the most informative variable features, a two-stage brute-force combinatorial strategy was employed. In the first stage, all combinations of variable features were evaluated at k-max = 4, where k-max denotes the number of variable features to be combined with the nine core features, yielding approximately 150,000 combinations across the 44 variable features. Features were ranked by their frequency of appearance in high-performing combinations, and the top 16 variable features were carried forward alongside the 9 core features, giving a reduced set of 25 features for the second stage.

In the second stage, the 25-feature set (9 core + 16 variable) was subjected to exhaustive brute-force combinatorial evaluation using both Random Forest and XGBoost as scoring algorithms. Given that the reduced variable feature set yields only 65,535 total combinations across all k-max values, this stage was computationally tractable in its entirety. A-EPAR values were used to identify the optimal feature subset for each algorithm, resulting in a final set of 17 features for Random Forest and 18 features for XGBoost.

#### Algorithm selection

The finalized feature sets were subsequently used to benchmark ten classification algorithms: XGBoost, LightGBM, Gradient Boosting, Random Forest, Support Vector Machine, Decision Tree, K-Nearest Neighbors, Logistic Regression, Linear Regression, and Naïve Bayes. Rather than averaging EPR across all thresholds, performance was visualized as a heatmap spanning deciles from 10% to 100%, enabling a comprehensive comparison of target recovery across algorithms and thresholds.

### Model training and evaluation

The 18-feature SMARTIE model was trained on TRIBE data from NonA experiments in *Drosophila* brain, with reanalyzed targets designated as positives (GSE116400). For each gene, the model outputs a probabilistic score between 0 and 1 reflecting the likelihood of it being a bona fide RBP target. Genes are ranked in decreasing order of probability to prioritize candidates for downstream validation.

The model was subsequentially tested on TRIBE data from additional RBPs obtained from previously published studies (Table 1). Supervised training was employed to learn nonlinear decision boundaries distinguishing target from non-target genes, with each gene assigned a probability score reflecting its likelihood of being a genuine target. Model performance was assessed using cross-validation, with predictive accuracy quantified by the area under the receiver operating characteristic curve (ROC-AUC). To quantify model performance throughout this process, a metric termed EPAR was defined. For each RBP, the top N SMARTIE-ranked predictions, where N corresponds to a given decile fraction of the reanalyzed target list, were intersected with the full reanalyzed target list, and the percentage overlap was computed. This was repeated across deciles from 10% to 100% to generate a comprehensive performance profile.

### Motif and metagene analysis

Motif analysis was conducted using the MEME suite, while metagene plots were generated using the Guitar R package.

## Tools and Software

**Table 3:**
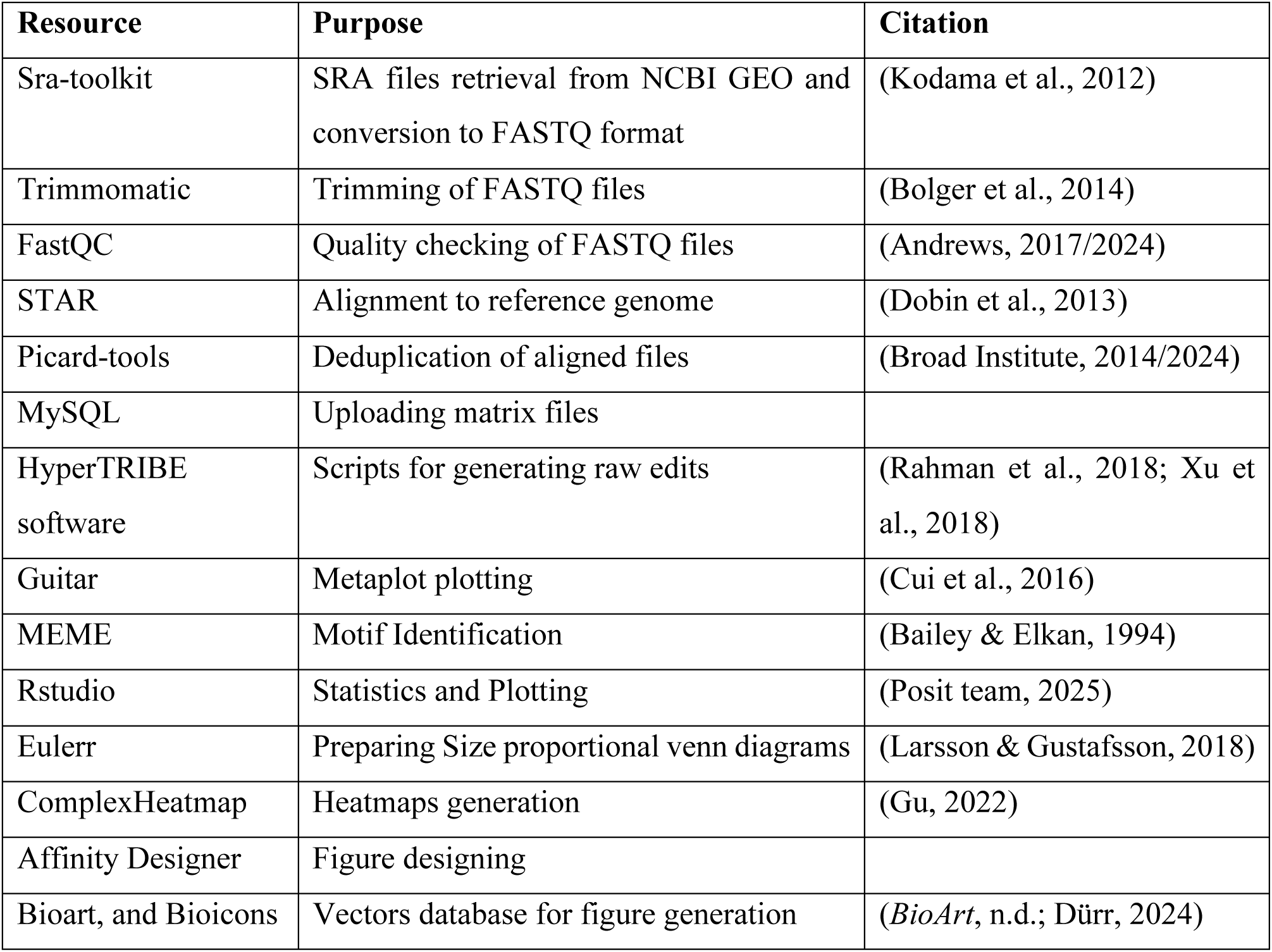
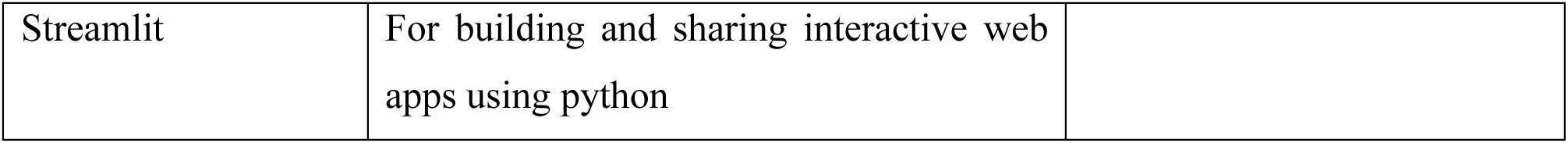
List of Tools and Software used in this study.

### Data availability

Detailed information about datasets downloaded for the study and benchmarking of SMARTIE is present in Table. S1. Reanalysed list of targets for the published datasets is available in Table. S2. For the list of 53 constructed features, divided based on their feature categories, please refer to Table. S3. SMARTIE predicted targets for two replicates and single replicates are provided in Table S4 and S5 respectively.

### Code availability

The SMARTIE package and code used for designing SMARTIE are available at the repository: https://github.com/toolsmnl/SMARTIE. In addition, a GUI based version of SMARTIE is also provided. SMARTIE can be customized by users based on their requirements. The code and instructions for the same are also available on the same repository.

## Acknowledgements

We thank Mani Ramaswami, Amanjot Singh and Arvind Reddy for their useful insights on the manuscript. We also thank Baskar Lab and the Param Himalaya Supercomputing Facility at IIT Mandi for the service and resources. Claude (Anthropic) was used to assist with script development, manuscript language editing, and graphical design. This work was supported by DBT / Wellcome Trust India Alliance (IA/I/19/1/504286) and DBT-GET (BT/PR38373/GET/119/330/2020) grants.

## Author Contributions

Conceptualization: OK and BB

Methodology: OK, UK, GA, RY and BB

Investigation: OK and UK

Funding Acquisition: BB

Project Administration: BB

Supervision: BB

Additional Supervision: RY and GA

Writing - original draft: OK and BB

Writing - review and editing: OK, UK, GA, RY and BB

## Ethical Declarations

The authors declare no competing interests

## Supplementary Information

Supplementary information (Table S1-5 and Figures. S1-6)

**Table. S1: Information of Datasets used for training and testing SMARTIE**

**Table. S2: List of targets identified by reanalysing the published datasets**

**Table. S3: List of all the 53 constructed features**

**Table. S4: List of targets ranked by SMARTIE**

**Table. S5: List of targets ranked by SMARTIE for single replicates**

**Figure. S1:**
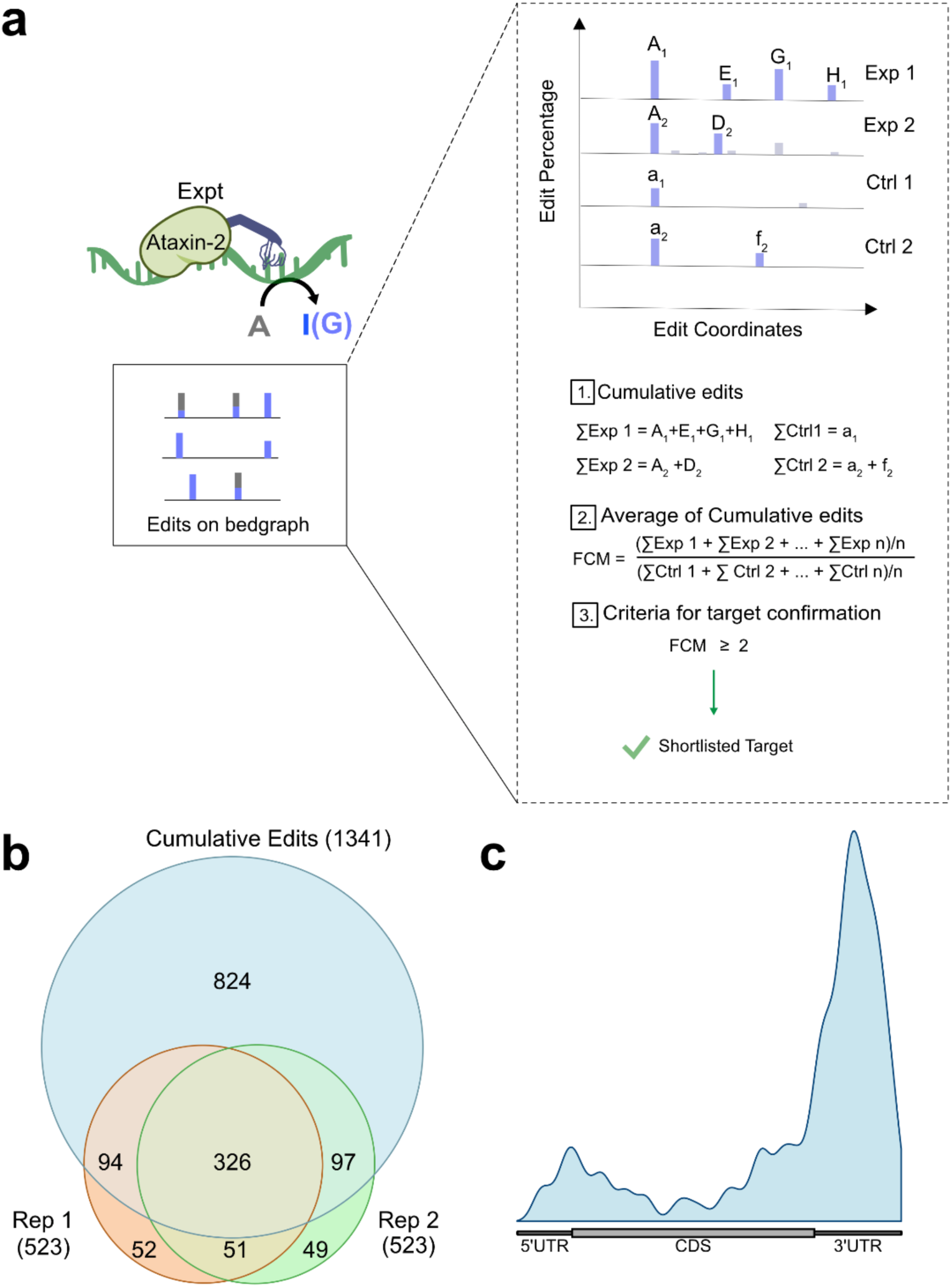
Cumulative transcript wide editing signature improves TRIBE target identification: (a) An illustration on TRIBE RNA editing by the ADARcd and the process of calculating cumulative edit percentage and deriving fold-change to identify potential RNA targets.(b) A Venn diagram illustrating the intersection of Ataxin-2 RNA targets discovered from individual replicates using conventional analysis and those identified using cumulative edit method. (c) Metaplot for Ataxin-2 RNA targets uniquely identified based on cumulative edits on transcripts.

**Figure. S2.**
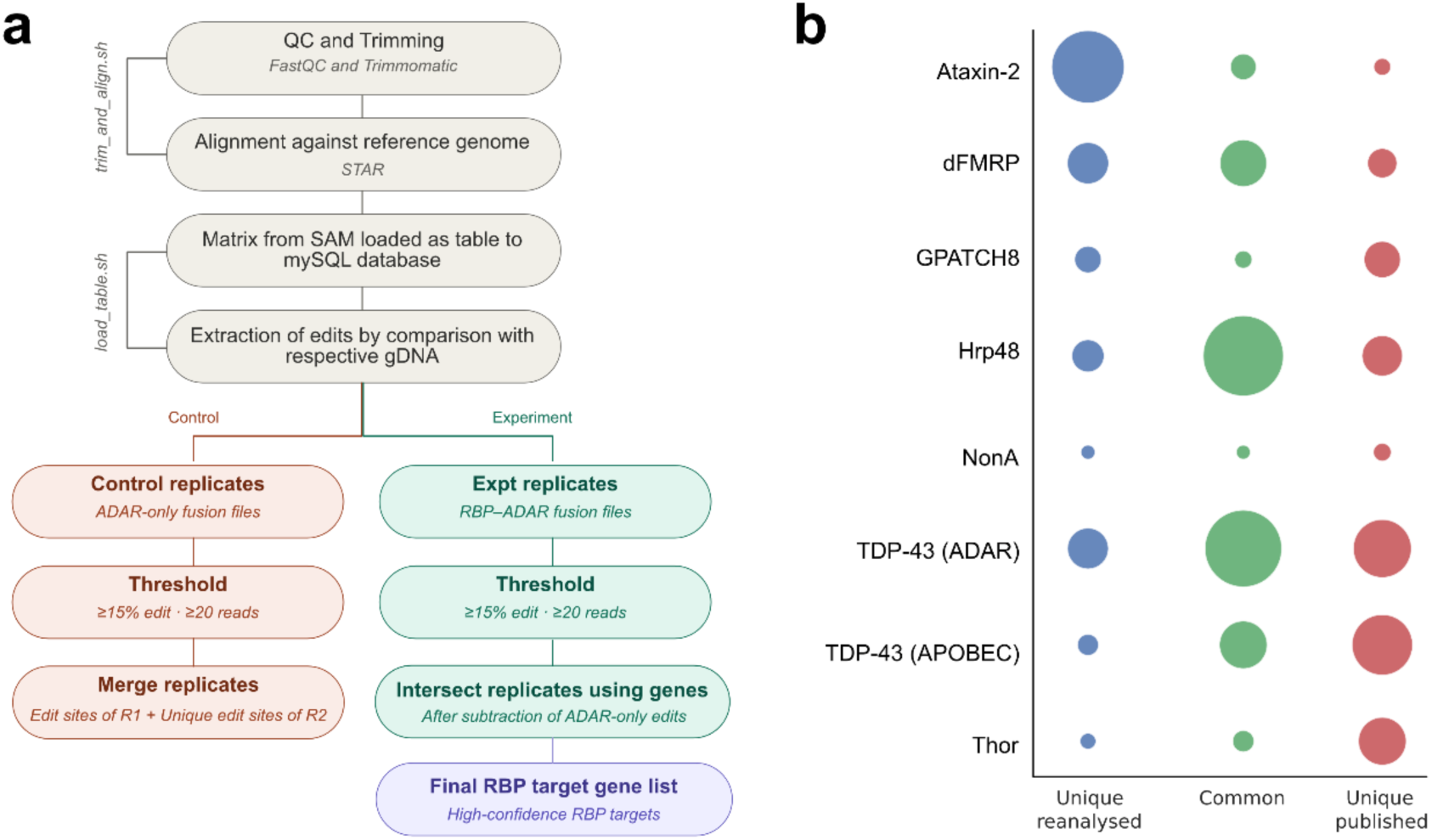
Reanalysis of published TRIBE datasets for RBPs used for training and testing machine-learning models. (a) Schematic of the reanalysis pipeline used to process publicly available TRIBE datasets and generate standardized target lists. (b) Dot plot showing the overlap between reanalyzed (blue) and published (red) target sets for each RBP, with genes common to both represented in green.

**Figure. S3:**
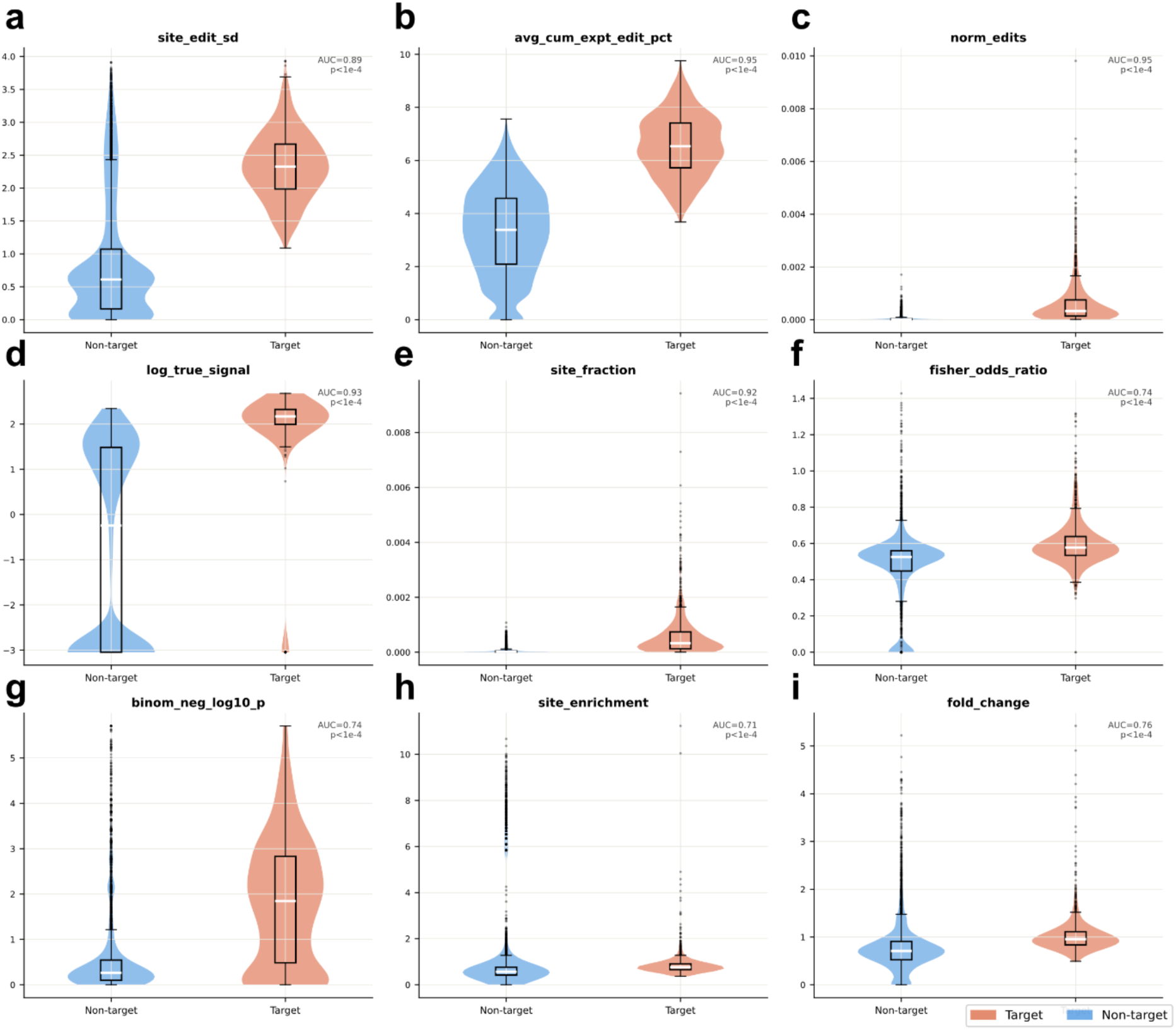
The features that distinguish experimental TRIBE data from control. Values for each of the biologically relevant core features were calculated for target and non-target genes and plotted.

**Figure. S4:**
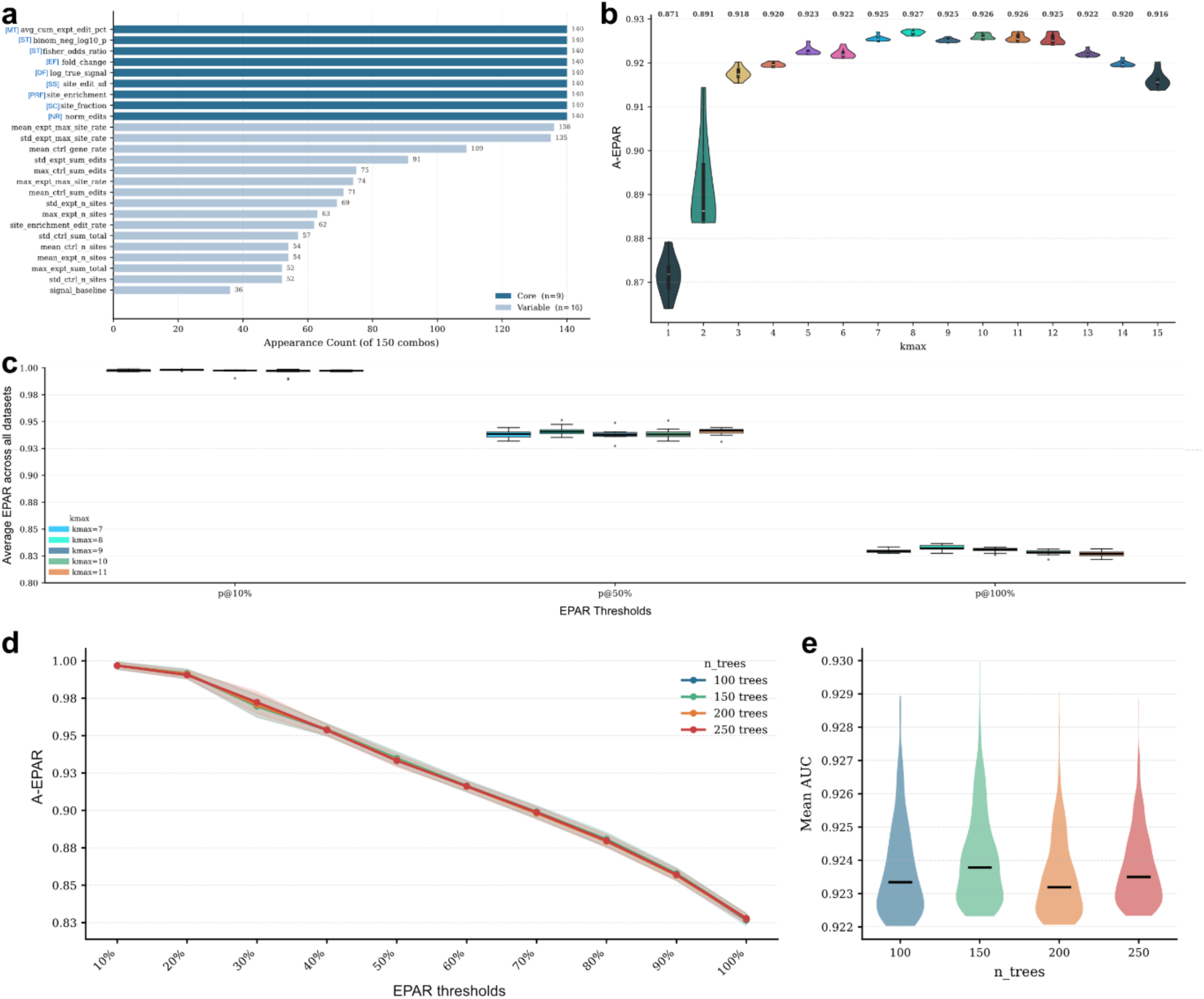
Evaluation of Random Forest algorithm for SMARTIE using brute force strategy. (a) Bar plot depicting the frequency of appearance of each feature across the top 10 combinations at each k-max value from 1 to 15, highlighting the most consistently informative features. (b) Violin plots showing the distribution of A-EPAR scores across the top 10 feature combinations at each k-max from 1 to 15, illustrating how overall model performance evolves with increasing feature set size. (c) Box plots comparing combination-level EPAR across various classification thresholds for k-max 7-11. (d and e) Effect of tree size on the ability of Random Forest to pick the features.

**Figure. S5:**
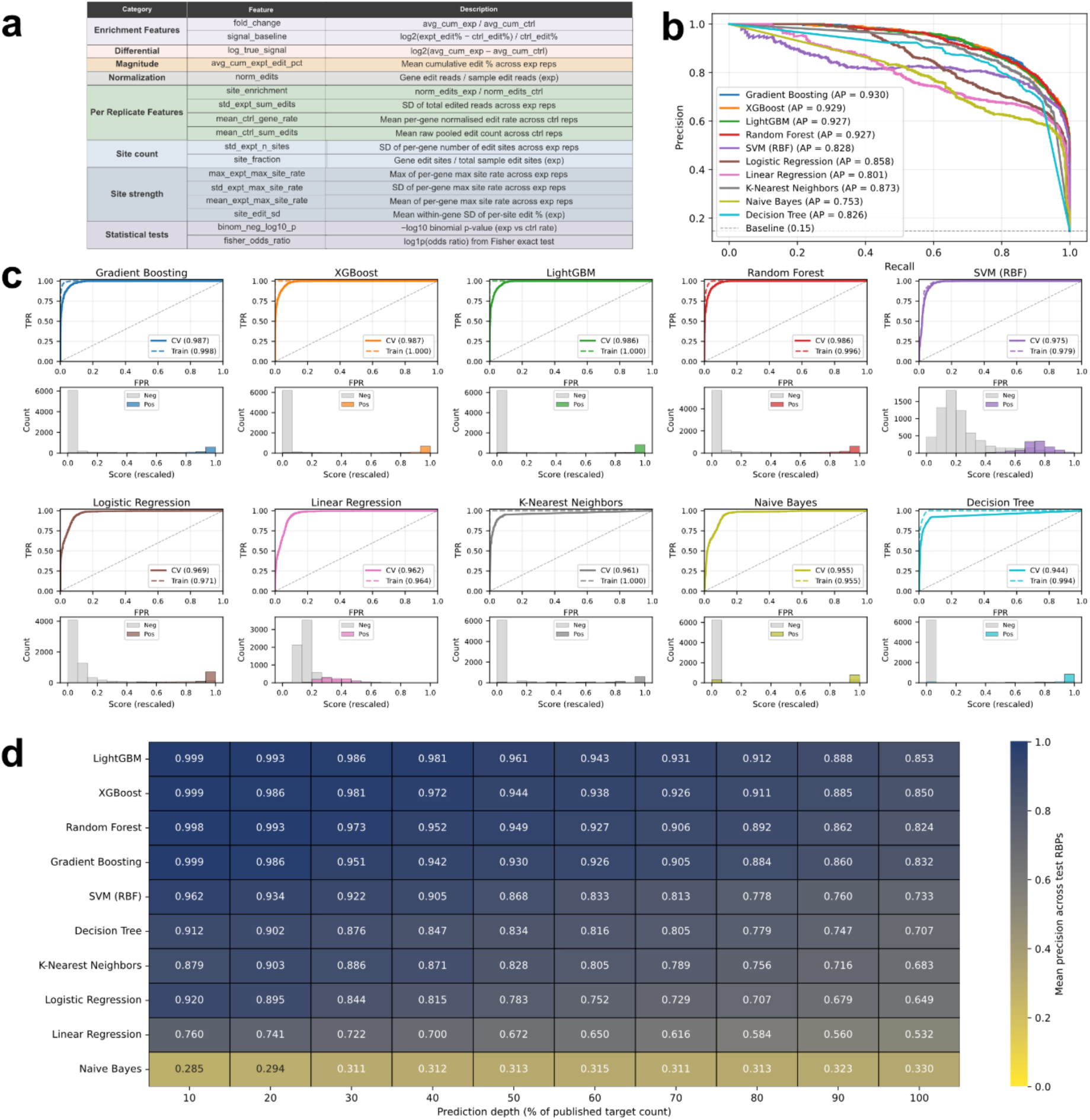
Comparative evaluation of classification models algorithms. (a) Description for each of the 17 features selected for training SMARTIE. (b) ROC curves for all ten candidate models evaluated on the training dataset, illustrating their ability to discriminate RBP targets from non-targets. (c) Paired comparison of training and cross-validation ROC curves (top) alongside the distribution of predicted scores for targets and non-targets (bottom), used jointly to assess model overfitting and discriminative capacity. (d) Heatmap depicting average EPAR at each decile threshold across all held-out testing datasets, enabling quantitative comparison of target recovery performance across models.

**Figure. S6:**
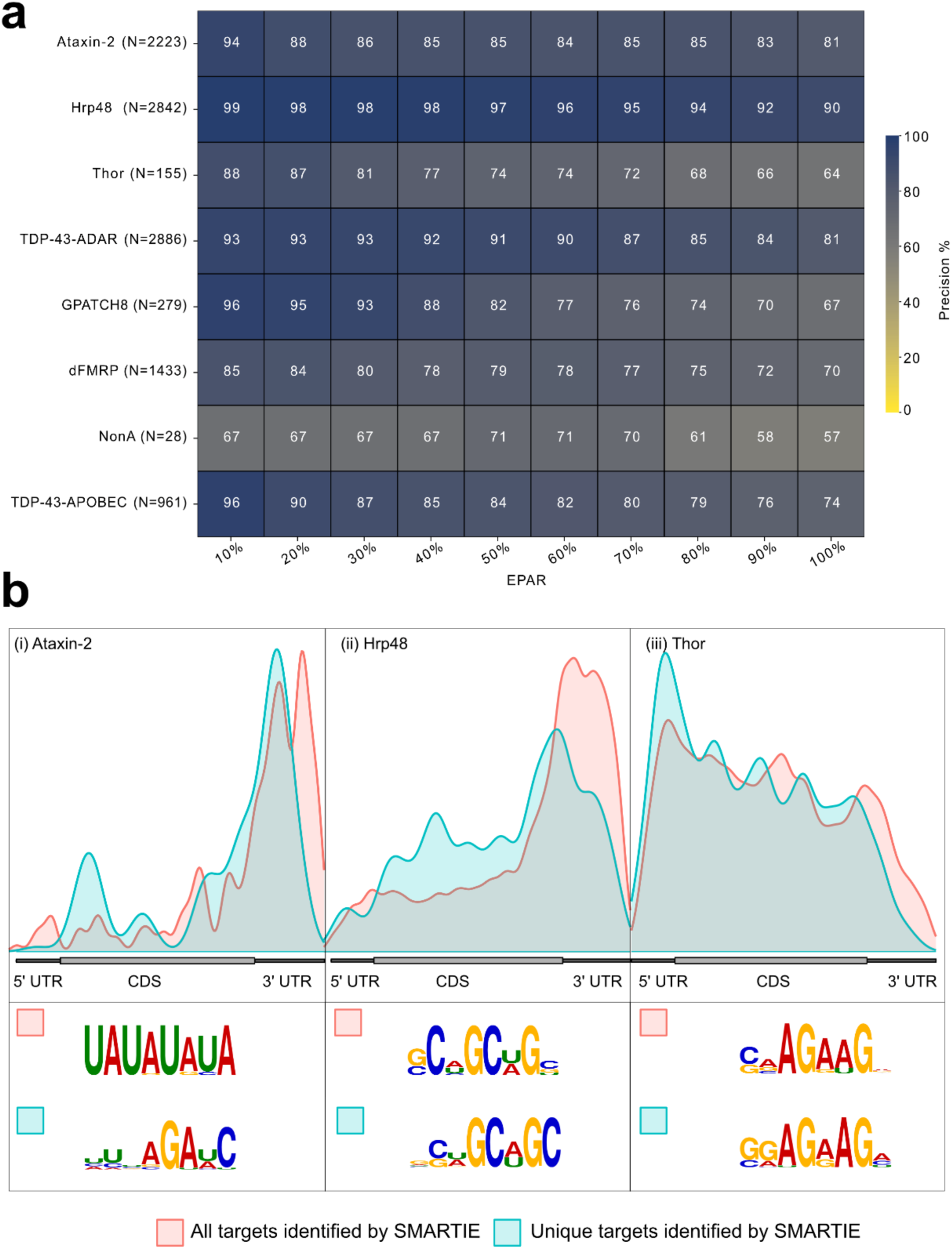
Benchmarking SMARTIE with single replicate using TRIBE and STAMP datasets. **(a)** A heatmap for decile-based analysis of SMARTIE-predicted target intersected with the reanalysed list of targets. (b) Metaplot and motifs for targets uniquely identified by SMARTIE using single replicate data.

**Figure S7:**
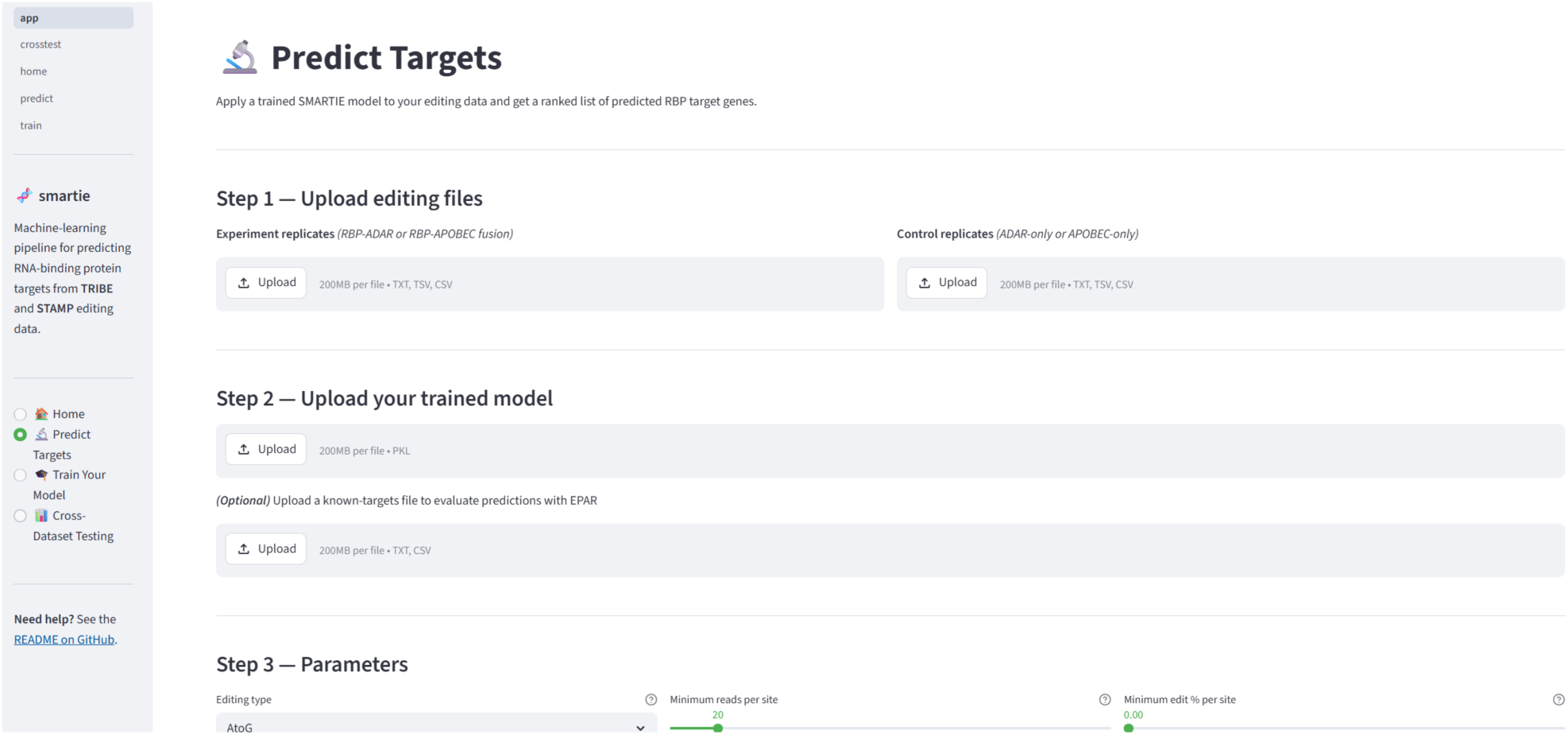
Graphical User interface for SMARTIE built using Streamlit. A screen short of SMARTIE’s GUI to generate downloadable ranked target predictions from raw experiment and control files.

