## Supplementary material for "SMARTIE: A Machine-Learning approach for investigating RBP-RNA interactions identified by Editing": Table S1

| Study | Sample | Designation | GEO ID | Reference |
| --- | --- | --- | --- | --- |
| Ataxin-2 TRIBE in fly neurons | Ataxin-2 WT TRIBE | Experiment | [GSE153985](https://www.ncbi.nlm.nih.gov/geo/query/acc.cgi?acc=GSE153985) | Singh et al, 2021 |
|  | UAS-Ataxin-2 WT | Background |  |  |
|  | Elav-GAL4 | Background |  |  |
|  | ADAR-only | Control | [GSE282990](https://www.ncbi.nlm.nih.gov/geo/query/acc.cgi?acc=GSE282990) | Koppaka et al, 2025 |
| NonA TRIBE in S2 Cells | NonA-ADAR | Experiment | [GSE78065](https://www.ncbi.nlm.nih.gov/geo/query/acc.cgi?acc=GSE78065) | McMahon et al, 2016 |
|  | ADAR only | Control |  |  |
| Hrp48 HyperTRIBE in S2 cells | Hrp48-HyperADAR | Experiment | [GSE102814](https://www.ncbi.nlm.nih.gov/geo/query/acc.cgi?acc=GSE102814) | Wu et al, 2018 |
|  | HyperADAR only | Control |  |  |
| Thor HyperTRIBE in S2 cells | Thor-HyperADAR | Experiment | [GSE153346](https://www.ncbi.nlm.nih.gov/geo/query/acc.cgi?acc=GSE153346) | Jin et al, 2020 |
|  | HyperADAR only | Control |  |  |
| TDP-43 HyperTRIBE and STAMP in HEK293T | TDP-43 HyperADAR | Experiment | [GSE223556](https://www.ncbi.nlm.nih.gov/geo/query/acc.cgi?acc=GSE223556) | Abruzzi et al, 2023 |
|  | HyperADAR only | Control |  |  |
|  | TDP-43 APOBEC | Experiment |  |  |
|  | APOBEC only | Control |  |  |
| dFMRP in S2 cells | dFMRP HyperADARcd | Experiment | [GSE102814](https://www.ncbi.nlm.nih.gov/geo/query/acc.cgi?acc=GSE102814) | Wu et al, 2018 |
|  | HyperADARcd only | Control |  |  |
| GPATCH8 in HEK293T | GPATCH8 ADARcd | Experiment | [GSE242093](https://www.ncbi.nlm.nih.gov/geo/query/acc.cgi?acc=GSE242093) | Benbarche et al, 2024 |
|  | ADARcd only | Control |  |  |
| NonA TRIBE in Brain | NonA ADARcd | Experiment | [GSE116400](https://www.ncbi.nlm.nih.gov/geo/query/acc.cgi?acc=GSE116400) | Unpublished |
|  | ADARcd only | Control |  |  |
